# Cytoplasmic streaming drifts the polarity cue and specifies the cell polarity in *Caenorhabditis elegans* zygotes

**DOI:** 10.1101/2019.12.23.887620

**Authors:** Kenji Kimura, Akatsuki Kimura

**Affiliations:** Cell Architecture Laboratory, Structural Biology Center, National Institute of Genetics, Mishima 411-8540, Japan; Department of Biological Science, Kwansei Gakuin University, 2-1 Gakuen, Sanda 669-1337, Japan; Department of Genetics, School of Life Science, The Graduate University for Advanced Studies, SOKENDAI, Mishima 411-8540, Japan

**Keywords:** sperm entry site, polarity, symmetry breaking, cytoplasmic streaming, *C. elegans*

## Abstract

Cell polarisation is required to define body axes during development. The position of spatial cues for polarisation is critical to direct the body axes. In *Caenorhabditis elegans* zygotes, the sperm-derived pronucleus/centrosome complex (SPCC) serves as the spatial cue to specify the anterior–posterior axis. Approximately 30 minutes after fertilisation, the contractility of the cell cortex is relaxed near the SPCC, which is the earliest sign of polarisation and called symmetry breaking (SB). It is unclear how the position of SPCC at SB is determined after fertilisation. Here, we show that SPCC drifts dynamically through the cell-wide flow of the cytoplasm, called meiotic cytoplasmic streaming. This flow occasionally brings SPCC to the opposite side of the sperm entry site before SB. Our results demonstrate that cytoplasmic flow determines stochastically the position of the spatial cue of the body axis, even in an organism like *C. elegans* for which development is stereotyped.

## Introduction

Cell polarisation is a fundamental step in animal development; asymmetric distribution of the determinants of cell fate within a cell establishes the cell polarity. Unidirectionally reinforced transport of polarity determinants through a well-organised cytoskeleton contributes in establishing the polarity in many cases^1-5^. Internal or external cues that induce cell polarisation such as the sperm entry site, the sperm-derived aster, or surrounding cells provide positional information for the proper orientation of embryonic polarity^6, 7^. The zygotes of *C. elegans* present a paradigm for polarisation in which the sperm-derived pronucleus/centrosome complex (SPCC) possess cues that establish the anterior–posterior (AP) polarity^8^.

In *C. elegans* zygotes, the position of SPCC at the time of “symmetry breaking (SB)” is known to determine the AP polarity. SB occurs approximately 30 minutes after fertilisation when local relaxation of contractility of the cell cortex in the vicinity of SPCC occurs, which is the earliest sign of the establishment of the AP polarity^9, 10^. At this time, Aurora A kinase (AIR-1) is known to relocate from the cytoplasm to the centrosomes^11-13^. The local relaxation triggers unidirectional advection of the cortical actomyosin that brings the anterior PAR proteins to the future anterior cortex, and simultaneously the posterior PAR proteins are allowed to localise to the future posterior cortex^1, 14^. In addition, a cytoplasmic flow is generated in the inner cytoplasm in a counter-direction of the cortical advection^15^, which potentially promotes the loading of the posterior PAR proteins to the future posterior cortex^16^. As a result, the side with SPCC becomes the posterior and the other side becomes the anterior. Thus, the position of SPCC at SB defines AP polarity in *C. elegans* zygotes.

How the position of the spatial cue (SPCC) at SB is determined is a critical question to understand the mechanism of polarity specification. The clear answer, however, has not been obtained yet. Because SPCC is derived from the sperm, the site of sperm entry is suspected to have a critical role. In fact, the site of sperm entry seems to correlate with the position of SPCC at SB, and thus with the future posterior pole^8^. In the *C. elegans* hermaphrodites, the sperm usually fuses with the leading edge of the oocyte, migrating into a sperm storage organ (spermatheca)^17^. We refer to the side of the leading edge as the proximal side, and the opposite side as the distal side (Fig. 1A). Because the oocyte nucleus is positioned at the distal side in oocytes, where the polar body extrusions occur after fertilisation, the distal side is marked by the position of the first polar body (Fig. 1A and 1B). In most cases, the proximal side becomes the posterior of the zygote. We call this “PP-type polarisation” (Fig. 1C). Importantly, the correlation between the proximal side of the oocyte and the posterior of the zygote is not perfect. In some cases, the distal side becomes the posterior (as in “DP-type polarisation”, Fig. 1C). DP-type polarisation was observed when a sperm-less hermaphrodite (*fem-1(hc17)* strain) was mated with wild-type males^8^. This treatment was considered to alter the position of sperm entry and increased the chance for the sperm to enter from the distal side of the oocyte. This scenario is based on two assumptions as follows: (i) the sperm can enter from both proximal and distal sides of the oocyte, and (ii) SPCC does not move much until SB after fertilisation. However, it was unclear whether the sperm actually entered from the distal side when DP-type polarisation occurred. The previous observation was based on Nomarski optics, and the position of sperm entry was judged by the position where the sperm pronucleus becomes visible, which is approximately 30 min after fertilisation. It is not clear as to how much the SPCC moves in the zygotes during the 30-min period after fertilisation. Therefore, it remains elusive how the position of SPCC at SB is determined.

**Figure 1.**
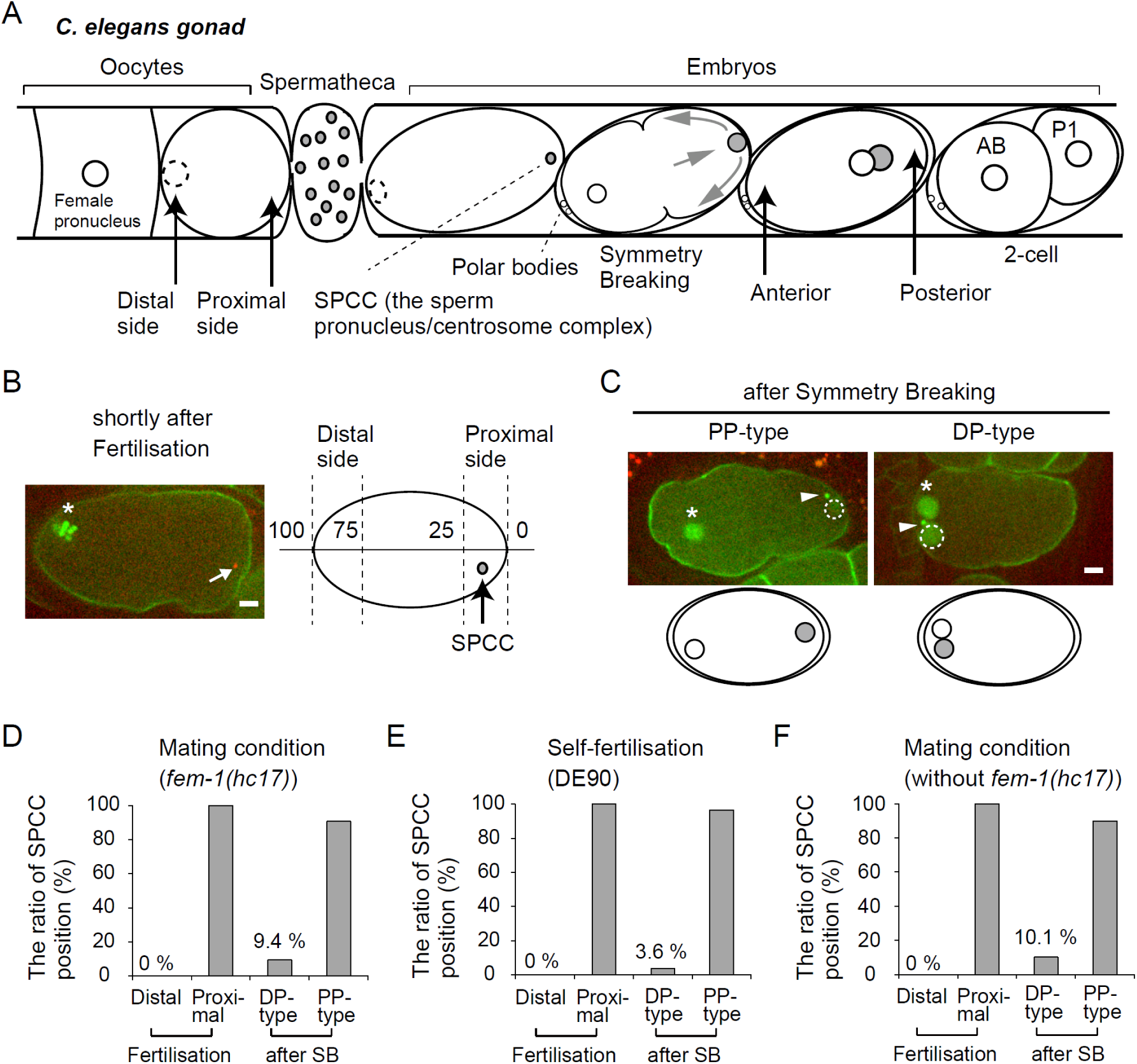
The sperm enters only through the proximal side of the oocyte. (A) Schematic of the *Caenorhabditis elegans* gonad and the definition of the proximal and distal sides. (B) Fluorescence confocal image of zygote expressing GFP::PH, GFP::histone and TBG-1::GFP in a *fer-1(hc17)* hermaphrodite crossed to a *him-5(e1490)* male expressing mCherry::histone at a time immediately after fertilisation. Asterisks indicate the oocyte-derived chromosomes. White arrow indicates the position of the sperm pronucleus/centrosome complex (SPCC). The distal quartile and proximal quartile of the oocyte or zygote before polarisation are defined as the distal and proximal sides, respectively, as shown in the right schematic. Scale bar, 5 μm. (C) Representative examples of the PP and DP-type after symmetry breaking (SB). Arrow heads indicate the centrosome. White dotted circles indicate the male pronucleus. Scale bar, 5 μm. (D-F) The ratios of the zygote of which SPCC is positioned in the distal or proximal side immediately after fertilisation and after SB under conditions of (D) mating using CAL0182 (*fer-1(hc17)* mutant) hermaphrodites and CAL0841 males, (E) self-fertilisation (DE90), and (F) mating using CAL1041 hermaphrodites and CAL1651 males are shown.

We suspected that the SPCC may move during the 30-min period. Before the appearance of sperm pronuclei, a collective flow of the cytoplasm called meiotic cytoplasmic streaming (MeiCS) occurs in *C. elegans* zygotes^18, 19^. We previously revealed the mechanism of the stochastic change of the flow direction of MeiCS^20^. In this study, we investigated the influence of MeiCS on the dynamics of SPCC before the onset of polarity establishment. Our extensive observations on the position of SPCC after fertilisation reveal that cytoplasmic streaming drastically influences the position of SPCC before SB and has a deterministic role in AP polarity.

## Results

### The Sperm Enters Only through the Proximal Side of the Oocyte

To clarify the position of sperm entry and of sperm-derived chromosomes after fertilisation, we conducted a time-lapse imaging of the zygote produced under the mating condition using the *fem-1(hc17)* strain (Fig. 1B), following the procedure described previously^8^. We visualised the chromosomes of the *fem-1(hc17)* oocyte by using a green fluorescent protein fused to the histone protein (GFP::histone) under the control of *pie-1* promoter, and visualised the sperm-derived chromosomes derived from a male by using a red fluorescent protein (mCherry::histone) under the control of *spe-11* promoter. To obtain males efficiently, we utilised the *him-5(e1490)* mutant that produces male progeny at an increased frequency^21^. Consistent with the results of the previous study^8^, we obtained zygotes with the DP-type polarisation at 9.4% frequency (5/53 embryos; Fig. 1D). More specifically, the sperm-derived chromosomes were positioned in the distal area (75–100% along the long axis of the zygote; Fig. 1B) at a timing known as “symmetry breaking (SB),” which is the timing of the earliest sign of polarity establishment (i.e. the local relaxation of cortical contractility) and is approximately 30 min after fertilisation. Unexpectedly, under the same mating condition as in the previous study^8^, all sperms entered from the proximal side (52 embryos); SPCC was found within the range of 0–25% in all cases immediately after fertilisation (Fig. 1D). This frequency (0/52) was significantly lower (*p* = 0.0058) than the frequency of observing the DP-type polarisation (5/53). Thus, our live-cell imaging indicates that the site of sperm entry is always the proximal side even in the case of DP-type polarisation.

To demonstrate that the DP-type polarisation is observed not only in the *fem-1* mutant, we also conducted experiments using non-mutant strains (“wild-type,” but encodes fluorescently-labelled proteins). The DP-type polarisation was observed both in the self-fertilisation and mating conditions [Fig. 1E and 1F; 3.6% (3/84) and 10.1% (7/69), respectively]. Importantly, under these conditions also, the site of sperm entry was always the proximal side [Fig. 1E (89 embryos), Fig. 1F (51 embryos)]. This result is the first clear demonstration that the sperm does not enter from the distal side of the oocyte and that the site of sperm entry does not specify the posterior pole in the *C. elegans* zygote.

### SPCC Occasionally Moves Away from the Sperm Entry Point after Fertilisation

The earliest sign of polarity establishment is the local relaxation of cortical contractility observed at the time of SB, which is ∼30 min after the fertilisation. The relaxation occurs at the nearest cortex of the centrosomes associated with SPCC^9, 10^. Consistent with this view, when the DP-type polarisation occurs, SPCC is observed in the distal side upon SB (Fig. 1C)^8^. In contrast, for the DP-type polarisation, the sperm entry site does not coincide with the posterior pole and the position of SPCC at SB. This disagreement inevitably suggested that SPCC moved from the sperm entry site (i.e. the proximal side) to the distal side for the DP-type polarisation. We succeeded in directly demonstrating that this is actually the case, by optimising the image acquisition condition to track the position of SPCC at short time intervals, with less photobleaching (Fig. 2A; Sup Movie 1). We quantified the position along the long axis of the zygote from the images (Fig. 2B). When the zygotes conduct the DP-type polarisation, SPCC starts from the proximal side and drifts to the distal side after fertilisation (Fig. 2A lower; Sup Movie 2; Fig. 2C, red symbols and lines). This observation indicates that the position of the spatial cue of polarity (SPCC) is stochastically changed from the sperm entry site before polarity establishment (SB).

**Figure 2.**
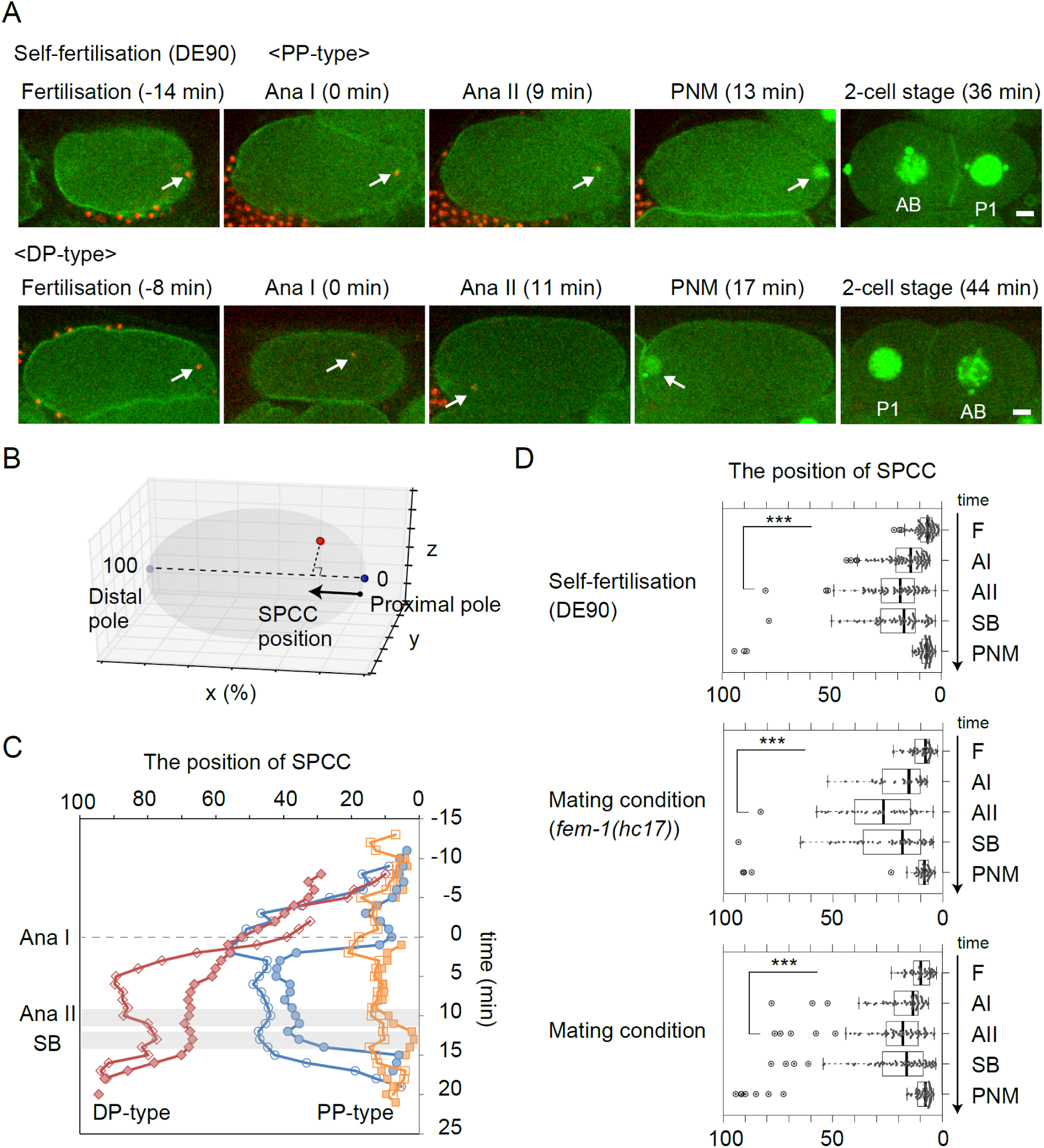
SPCC stochastically moves within the zygote before cell polarisation. (A) Time series images of *Caenorhabditis elegans* zygotes (DE90 strain) expressing mCherry::histone, GFP::PH, GFP::histone and TBG-1::GFP at fertilisation, meiotic anaphase I (Ana I), meiotic anaphase II (Ana II), before pronuclear migration (PNM) and 2-cell stage under the condition of self-fertilisation. White arrows indicate the position of the sperm pronucleus/centrosome complex (SPCC). Representative examples of the DP-type are shown in the bottom panels. The timing of meiotic anaphase I is set to 0 min. Scale bars, 5 μm. (B) The position of SPCC along the long axis of the zygote is evaluated as shown in the diagram and is plotted in (C). The proximal pole and the distal pole are set to 0 and 100, respectively. (C) Six representative examples of SPCC dynamics are shown. For details see the text. (D) Box plots of the position of SPCC along the long axis of the zygote at the timing of fertilisation (F), meiotic anaphase I (AI), meiotic anaphase II (AII), immediately after symmetry breaking (SB) and before pronuclear migration (PNM) under each condition. Asterisks indicate significant differences between F and AI (∗∗∗*P <0.001*).

Another interesting observation is that, even for the PP-type polarisation in which the sperm entry site coincides with the posterior pole, some SPCC moved dynamically to an almost indistinguishable level that causes the DP-type polarisation until meiotic anaphase II (Fig. 2C, blue symbols and lines). We evaluated the distribution of the position of SPCC at the time of fertilisation, meiotic anaphase I, meiotic anaphase II, SB, and just before pronuclear migration from all our tracking data (Fig. 2D). The quantification revealed that a subset of SPCC (15% and 23% under the self-fertilisation and mating conditions, respectively) was positioned beyond one-quarter of the long axis from the proximal pole (x-position ≥ 25%, Fig. 2B) at meiotic anaphase I, which is approximately 10 min after fertilisation (Fig. 2D), whereas all the SPCCs were found within the proximal side (0–25%), immediately after fertilisation (Fig. 2D). Some of them further moved toward the distal side and the distribution of the position of SPCC at symmetry breaking had a wide range (Fig. 2D). In contrast, a majority of SPCCs were positioned within one-quarter of the long axis (x-position < 25%) throughout the period from fertilisation to the onset of male pronuclear migration (61% and 68% under the self-fertilisation and mating conditions, respectively; two examples are shown in the orange plots in Fig. 2C). In summary, based on the results of tracking analyses, we provide direct evidence that SPCC occasionally moves from the proximal side to the distal side to induce the DP-type polarisation. The analyses also suggested that we cannot predict the polarity until at least anaphase I, as the movements of SPCC for the PP-type and DP-type polarisation are indistinguishable before anaphase I. We will discuss the latter point at a later instance.

### Meiotic Cytoplasmic Streaming Moves the Sperm Complex before SB

It is a pertinent question as to what is driving the dynamic movement of SPCC within the zygote after fertilisation. In the *C. elegans* zygote, immediately after fertilisation and before SB, a global flow of the cytoplasm called as meiotic cytoplasmic streaming (MeiCS) is intermittently generated in a kinesin-1 (*unc-116*)-dependent manner^18-20^. We previously deciphered the mechanism of the stochastic emergence and reversal of the flow^20^. Because the timing of the movement of SPCC is consistent with that of MeiCS, we hypothesised that the drift of SPCC depends on MeiCS. Here, we provide two lines of evidence supporting this hypothesis. Firstly, we asked whether the direction of MeiCS is consistent with that of the movement of SPCC. We simultaneously observed SPCC and MeiCS using GFP-labelled yolk granules. We found that SPCC moved together with a flow of yolk granules (Fig. 3A and Sup Movie 3). A correlation was found in the direction between SPCC movement and a flow of peripheral yolk granules (*r* = 0.48, Fig. 3B). The result supports the idea that MeiCS drives the movement of SPCC before SB. Secondly, we determined whether the movement of SPCC was suppressed when we attenuated the MeiCS. In *unc-116*/kinesin-1 (RNAi) zygotes, the position of SPCC did not change from the proximal side (Fig. 3C and D). To exclude the possibility that kinesin-1 directly transports SPCC, we also inhibited MeiCS by knockdown of *yop-1; ret-1* genes encoding reticulon proteins are required for the integrity of the ER^22^. We previously showed that we can impair the collectivity of MeiCS by fragmenting the ER through *yop-1;ret-1* (RNAi), without inhibiting the motor activity of kinesin-1^20^. As in *unc-116*/kinesin-1 (RNAi) zygotes, the position of SPCC was restricted close to the proximal pole in the *yop-1; ret-1* (RNAi) zygotes (Fig. 3C). As a negative control, we impaired the actin-dependent cortical and cytoplasmic flow upon SB by knocking down myosin II. In a previous study, it was demonstrated that MeiCS does not require the actin cytoskeleton^18^. Under this condition, SPCC moved away from the proximal pole before meiotic anaphase II (Fig. 3C). These results indicate that MeiCS moves SPCCs from the entry point before SB that sometime causes the DP-type polarisation. The DP-type polarisation was never observed when MeiCS was impaired (n = 20 in *unc-116* (RNAi) zygotes and n = 23 in *yop-1;ret-1* (RNAi) zygotes).

**Figure 3.**
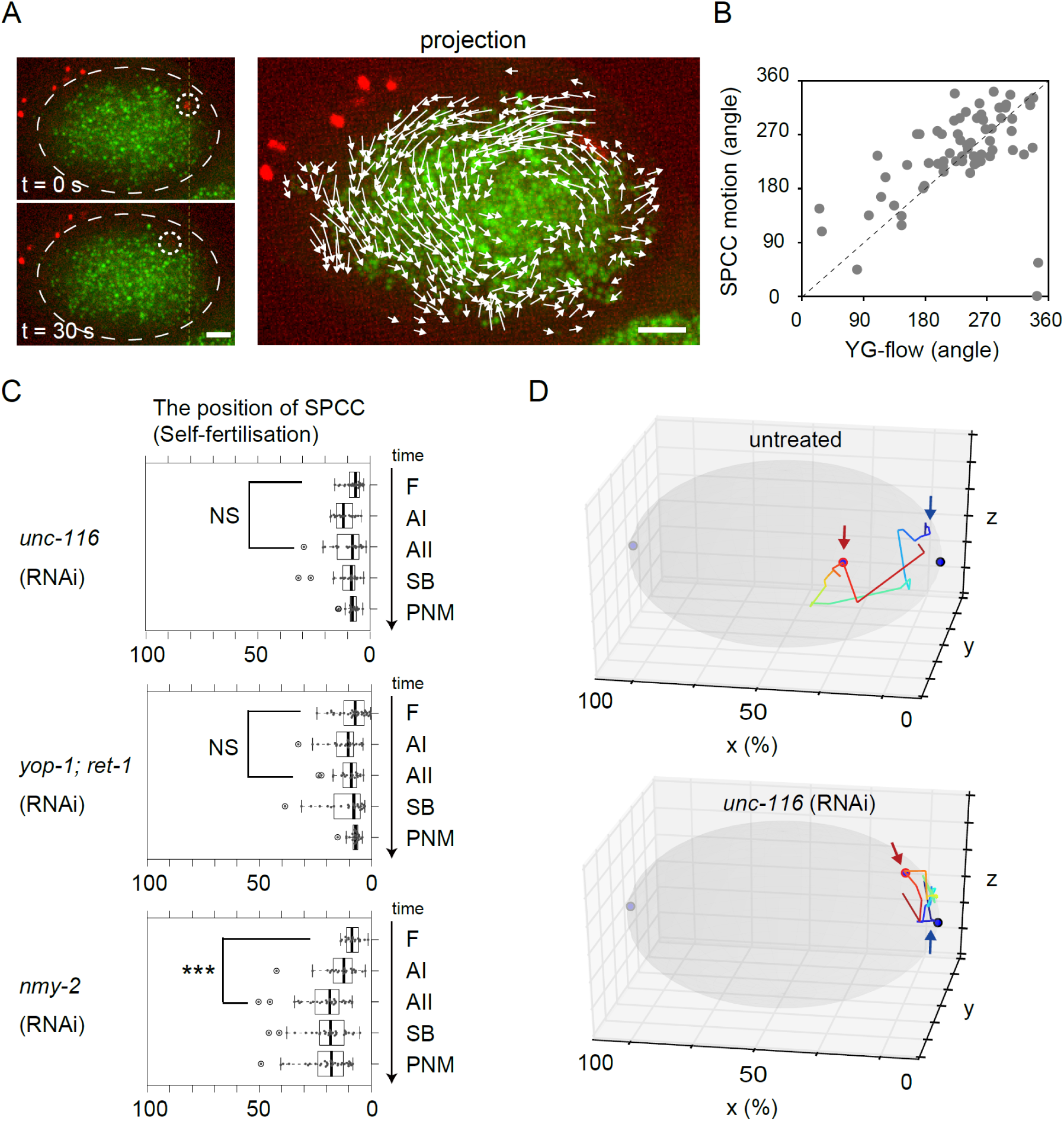
Meiotic cytoplasmic streaming moves sperm pronucleus/centrosome complex (SPCC). (A) The left two panels show time series of *Caenorhabditis elegans* zygote labelled yolk granules (VIT-2::GFP) and SPCC (mCherry::histone). The shape of the zygote is outlined by white dotted line. White dotted circles indicate SPCC. The right panel shows projections of sequential images of the zygote during MeiCS (six frames, time interval: 5 sec). White arrows indicate flow vectors of yolk granules (>1 μm). The yellow dotted line indicates the position of SPCC at 0 s. Scale bars, 5 μm. (B) Scatter plots of the angle (degree) of MeiCS and the direction of SPCC motion (69 frames from 5 zygotes). The mean angle of flow vectors of yolk granules within 7 μm radius of SPCC is calculated as the angle of MeiCS. (C) Box plots of the position of SPCC along the long axis of the zygote under conditions of *unc-116* RNAi, *yop-1; ret-1* RNAi and *nmy-2* RNAi) are shown as in Fig. 2D. Asterisks indicate significant differences between F and AI (∗∗∗*P <0.001*). NS, not significant. (D) Representative trajectory of SPCC from fertilisation to before pronuclear migration in untreated and *unc-116* (RNAi) zygote is shown by the coloured line. The grey ellipse represents the zygote and the blue and light blue circle indicates the proximal and distal poles, respectively. The blue arrows indicate the position of fertilisation. The red circles indicate the position of SPCC at the time of symmetry breaking.

### SPCC Moves Until Immediately Before SB, and Polarity Cannot be Predicted until the Timing of SB

At the timing of SB, the earliest sign of polarity (i.e. the local relaxation of cortical contractility) is observed^9, 10^. The question is when is the earliest timing to predict the future posterior pole (i.e. distal or proximal of the zygote). The prediction of the posterior may be made earlier than the actual sign of appearance of polarity at SB. We demonstrated that it is not upon the sperm entry, but later (Figs. 1 and 2). MeiCS that moves SPCC intensively (Fig. 3) is usually diminished before meiotic anaphase II, suggesting the possibility that SPCC does not move after meiotic anaphase II and that the position of SPCC at meiotic anaphase II practically determines polarity. To find the earliest-possible timing of the prediction of polarisation, we assembled all our tracking data. At SB, the most-distally-positioned SPCC of the PP-type polarisation was located more proximally (51.5%) than the most-proximally-positioned one of the DP-type polarisation (53.1%) (Fig. 4A and B). This result is consistent with the existing view that we can predict from the position of SPCC at SB. When we conduct this comparison at the meiotic anaphase II, which is a one-step earlier landmark that we set, the most-distally-positioned one of the PP-type polarisation was located more distally (55.2%) than the most-proximally-positioned one of the DP-type polarisation (43.9%) (Fig. 4B). This quantification indicates that we cannot predict the posterior from the position of SPCC at meiotic anaphase II. SPCCs are still moving to some extent after meiotic anaphase II, although the MeiCS is undetectable, and the movement can affect the decision of polarity. In conclusion, we demonstrated that SPCC moves until immediately before SB and clarified that we cannot predict the posterior pole from the position of SPCC until SB, when the final decision of polarity is made (Fig. 4C).

**Figure 4.**
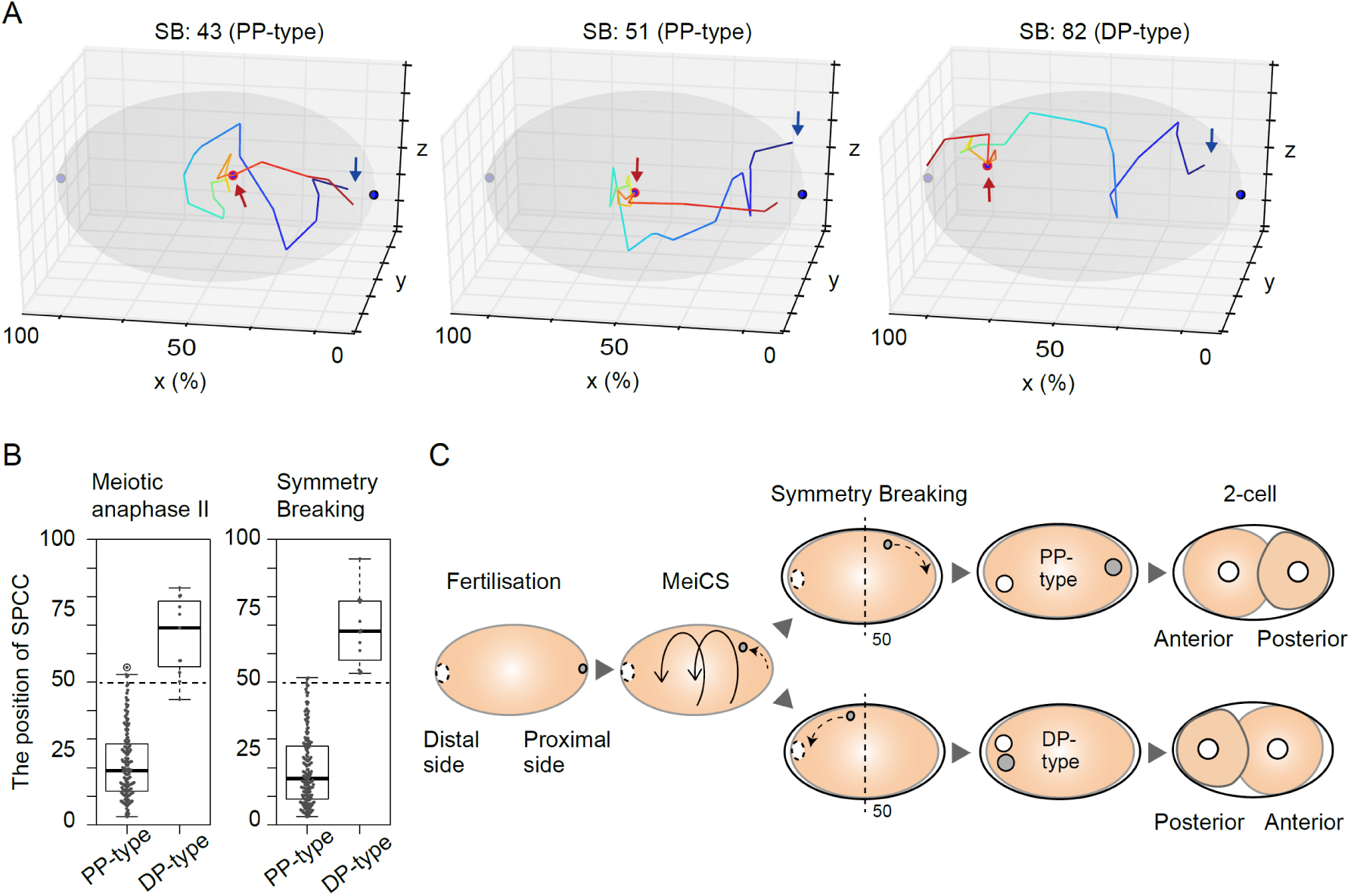
The sperm pronucleus/centrosome complex (SPCC) moves until immediately before symmetry breaking. (A) Trajectories of SPCC in the zygote in which symmetry breaking occurred when SPCC was positioned at 43, 51, and 82% of the long axis are shown as in Fig. 3D. (B) Box plots of the position of SPCC along the long axis of the zygote at meiotic anaphase II and symmetry breaking in the PP- and DP-type are shown as in Fig. 2D. (C) Summary of the SPCC dynamics during cell polarisation. For details see the text.

## Discussion

In this study, we revealed how the position of SPCC is determined after fertilisation before the onset of SB in the *C. elegans* zygote. We found that the sperm always enter from the proximal side of the oocyte. MeiCS drifts SPCC, and drift occasionally (∼10%) induces DP-type polarisation as a result. We show that SPCC moves more or less until immediately before SB, and thus, polarity cannot be predicted until SB, and not at anaphase II or earlier. The present study indicates that MeiCS has a critical role in the determination of cell polarity (PP-type or DP-type) and that the position of the spatial cue is stochastically determined depending on the stochastic nature of MeiCS before the zygotes proceed to a stereotyped pattern of development.

Is there a benefit for the cell to move SPCC with MeiCS? We think the movement of SPCC is a side effect of MeiCS, and possibly an unfavourable one. Our previous study suggested that a primary role of MeiCS is to increase the mobility of cortical granules for their efficient exocytosis at the meiotic anaphase I^20^. In contrast, the movement of SPCC is considered unfavourable for the cell as it may induce a contact between the oocyte chromosome and sperm chromosome before the oocyte meiosis^23^. The nature of MeiCS is likely designed to move cortical granules efficiently while keeping the movement of SPCC minimum. The flow direction of MeiCS is biased along the short axes of the zygote and, thus, it does not effectively moves SPCC along the long axis^20^. In addition, MeiCS is not stable and stochastically changes the direction^20^, which also makes a long-distance movement difficult. In fact, in the majority of zygotes (∼70%), SPCC stayed within 0–25% proximal side until SB.

Another mechanism to suppress the movement of SPCC by MeiCS is the anchoring of SPCC to the cell cortex^23^. Panzica et al. reported that, without the anchoring mechanism, SPCC detaches from the cortex and moves dynamically inside the cell^23^. Therefore, the movement of SPCC is likely a result of competition between the stirring effect of MeiCS and the anchoring mechanism. Our present results indicate that the MeiCS overcomes the anchoring mechanism to a certain extent. Even in those cases where the sperm chromosome gets closer to the oocyte chromosomes during the meiosis, we did not observe abnormal meiosis. It has been reported that, in the DP-type polarisation, where sperm chromosome should be near to the oocyte chromosomes before meiosis, the embryogenesis is normal^8^. Importantly, McNally et al. reported that the depletion of kinesin-1 subunit (KCA-1) promotes premature growth of sperm aster that can occasionally capture the oocyte chromosomes and causes a failure of meiosis^24^. These results collectively indicate a dual role of the kinesin-1 motor. Kinesin-1 generates MeiCS for efficient cortical granule exocytosis that stochastically leads the DP-polarisation. At the same time, kinesin-1 protects sperm chromosomes from mixing with oocyte chromosomes even when they are positioned in the vicinity. These mechanisms should make the mixture of cytoplasm and the integrity of chromosomes compatible. Considering the potential risk of MeiCS moving SPCC, there may be an undiscovered, greater benefit for the cell to conduct MeiCS.

Sperm-derived material provides a polarity cue to a fertilised zygote in some other species besides *C. elegans*. In the ascidian and frog egg, sperm entry or sperm-derived material affects the direction of cortical rotation, which polarises dorsal determinants toward the opposite side from sperm to specify the dorsal-ventral axis^5, 25, 26^. Massive cytoplasmic streaming often occurs after fertilisation in many species. In mouse, the site of sperm entry forms an actomyosin-rich cortex called the fertilisation cone that generates cytoplasmic flow through rhythmic contractions^27^. The movement of the sperm-derived materials after the fertilisation driven by cytoplasmic streaming might be a general phenomenon, that may affect the polarisation of the zygotes.

## Methods

### Strains and Manipulation of *C. elegans*

*C. elegans* strains were maintained using the standard techniques^28^. RNAi was performed by injecting double-stranded RNAs, as described previously^29^. For the mating experiments, CAL0182 (carrying *fem-1(hc17)*) hermaphrodites^30^ were grown at restrictive temperature (25 °C) and were mated to CAL0841 (carrying *him-5(e1490)*) males. CAL1041 hermaphrodites were mated to CAL1651 (carrying *him-5(e1490)*) males at 22 °C. For the analysis of the SPCC dynamics under self-fertilisation condition, DE90 strain was used^31^. For the analysis of cytoplasmic streaming, DH1033 strain was used^32^. The strains used in this study are listed in Table S1.

### Microscopy

Microscopic analysis was performed as described previously^29^. Briefly, worms were anesthetised with 1 mM levamisole and mounted on an agar pad. For the tracking analysis of the sperm complex within the zygote, eight or nine different focal planes (Z-interval = 3 µm) were acquired every 30 or 60 seconds. For the analysis of the flow field of MeiCS, three different focal planes (Z-interval = 3 µm) were acquired every 5 seconds.

### Image Analysis

For tracking analysis of SPCC, the movements of *spe-11p*::mCherry::histone signal, the male chromosomal marker, was manually tracked from time-lapse confocal images of the zygotes using Fiji software. We set the time of the onset of meiotic chromosomal separation as meiotic anaphase I or II, the time SPCC starts to move linearly to the cortex as the onset of SB, and the time just before male pronuclear migration as PNM (Fig. 2D). For drawing the trajectories of SPCC, we used Matplotlib (Line3DCollection) in a Python library.

For detection of the movements of yolk granules during MeiCS, optical flow analysis was conducted on time-lapse confocal fluorescence images of *C. elegans* zygotes. We used the Farnebäck method implemented in the OpenCV library. All images were rotated so that the long axis of the zygote was parallel to the horizontal axis before the optical flow analysis. To determine the correlation between the SPCC movement and the angle of MeiCS (Fig. 3B), the mean angle of the flow of yolk granules within 7 µm from SPCC was analysed for the period SPCC continuously moved on a single focal plane.

### Statistical Analysis

The probability that sperm enters the distal side was analysed using the BINOMDIST function (Microsoft Excel). The Student’s *t*-test was used to analyse the statistical difference between the SPCC distribution at fertilisation and at meiotic anaphase I (Fig. 2D and Fig. 3C). A Pearson correlation analysis was used to analyse the correlation between the SPCC movement and the flow direction of peripheral yolk granules (Fig. 3B).

### Data availability

The data supporting the findings of this study are available from the corresponding author upon request.

## Acknowledgments

Some strains were provided by the CGC, which is funded by NIH Office of Research Infrastructure Programs (P40 OD010440). We thank H. Sawa (National Institute of Genetics, Mishima, Japan), F. Motegi (National University of Singapore), and the members of cell architecture laboratory (National Institute of Genetics, Mishima, Japan) for their valuable comments and support.

## Author contributions

K.K. conceived the study. K.K. performed all the experiments. K.K. and A.K. analysed data and wrote the manuscript.

## References

1. Munro, E., Nance, J. & Priess, J.R. Cortical flows powered by asymmetrical contraction transport PAR proteins to establish and maintain anterior-posterior polarity in the early *C. elegans* embryo. Dev. Cell 7, 413–424 (2004).

2. Parton, R.M. et al. A PAR-1-dependent orientation gradient of dynamic microtubules directs posterior cargo transport in the *Drosophila* oocyte. J. Cell Biol. 194, 121–135 (2011).

3. Prodon, F., Sardet, C. & Nishida, H. Cortical and cytoplasmic flows driven by actin microfilaments polarize the cortical ER-mRNA domain along the a-v axis in ascidian oocytes. Dev. Biol. 313, 682–699 (2008).

4. Sardet, C., Paix, A., Prodon, F., Dru, P. & Chenevert, J. From oocyte to 16-cell stage: cytoplasmic and cortical reorganizations that pattern the ascidian embryo. Dev. Dyn. 236, 1716–1731 (2007).

5. Weaver, C. & Kimelman, D. Move it or lose it: axis specification in *Xenopus*. Development 131, 3491–3499 (2004).

6. Ajduk, A. & Zernicka-Goetz, M. Polarity and cell division orientation in the cleavage embryo: from worm to human. Mol. Hum. Reprod. 22, 691–703 (2016).

7. Chen, Q., Shi, J., Tao, Y. & Zernicka-Goetz, M. Tracing the origin of heterogeneity and symmetry breaking in the early mammalian embryo. Nature Commun. 9, 1819 (2018).

8. Goldstein, B. & Hird, S.N. Specification of the anteroposterior axis in *Caenorhabditis elegans*. Development 122, 1467–1474 (1996).

9. Bienkowska, D. & Cowan, C.R. Centrosomes can initiate a polarity axis from any position within one-cell *C. elegans* embryos. Curr. Biol. 22, 583–589 (2012).

10. O’Connell, K.F., Maxwell, K.N. & White, J.G. The *spd-2* gene is required for polarization of the anteroposterior axis and formation of the sperm asters in the *Caenorhabditis elegans* zygote. Dev. Biol. 222, 55–70 (2000).

11. Klinkert, K. et al. Aurora A depletion reveals centrosome-independent polarization mechanism in *Caenorhabditis elegans*. Elife 8 (2019).

12. Reich, J.D. et al. Regulated activation of the PAR polarity network ensures a timely and specific response to spatial cues. Curr. Biol. 29, 1911–1923. e1915 (2019).

13. Zhao, P. et al. Aurora-A breaks symmetry in contractile actomyosin networks independently of its role in centrosome maturation. Dev. Cell 48, 631–645. (2019).

14. Motegi, F. & Sugimoto, A. Sequential functioning of the ECT-2 RhoGEF, RHO-1 and CDC-42 establishes cell polarity in Caenorhabditis elegans embryos. Nat. Cell Biol. 8, 978–985 (2006).

15. Niwayama, R., Shinohara, K. & Kimura, A. Hydrodynamic property of the cytoplasm is sufficient to mediate cytoplasmic streaming in the *Caenorhabditis elegans* embryo. Proc. Natl. Acad. Sci. USA 108, 11900–11905 (2011).

16. Mittasch, M. et al. Non-invasive perturbations of intracellular flow reveal physical principles of cell organization. Nat. Cell Biol. 20, 344–351 (2018).

17. Samuel, A.D., Murthy, V.N. & Hengartner, M.O. Calcium dynamics during fertilization in *C. elegans*. BMC developmental biology 1, 8 (2001).

18. McNally, K.L., Martin, J.L., Ellefson, M. & McNally, F.J. Kinesin-dependent transport results in polarized migration of the nucleus in oocytes and inward movement of yolk granules in meiotic embryos. Dev. Biol. 339, 126–140 (2010).

19. Yang, H.Y., McNally, K. & McNally, F.J. MEI-1/katanin is required for translocation of the meiosis I spindle to the oocyte cortex in *C. elegans*. Dev. Biol. 260, 245–259 (2003).

20. Kimura, K. et al. Endoplasmic-reticulum-mediated microtubule alignment governs cytoplasmic streaming. Nat. Cell Biol. 19, 399–406 (2017).

21. Hodgkin, J., Horvitz, H.R. & Brenner, S. Nondisjunction mutants of the nematode *Caenorhabditis elegans*. Genetics 91, 67–94 (1979).

22. Audhya, A., Desai, A. & Oegema, K. A role for Rab5 in structuring the endoplasmic reticulum. J. Cell Biol. 178, 43–56 (2007).

23. Panzica, M.T., Marin, H.C., Reymann, A.C. & McNally, F.J. F-actin prevents interaction between sperm DNA and the oocyte meiotic spindle in *C. elegans*. J. Cell Biol. 216, 2273–2282 (2017).

24. McNally, K.L. et al. Kinesin-1 prevents capture of the oocyte meiotic spindle by the sperm aster. Dev. Cell 22, 788–798 (2012).

25. Roegiers, F., McDougall, A. & Sardet, C. The sperm entry point defines the orientation of the calcium-induced contraction wave that directs the first phase of cytoplasmic reorganization in the ascidian egg. Development 121, 3457–3466 (1995).

26. Houston, D.W. Cortical rotation and messenger RNA localization in *Xenopus* axis formation. Wiley Interdiscip Rev Dev Biol 1, 371–388 (2012).

27. Ajduk, A. et al. Rhythmic actomyosin-driven contractions induced by sperm entry predict mammalian embryo viability. Nature Commun. 2, 417 (2011).

28. Brenner, S. The genetics of *Caenorhabditis elegans*. Genetics 77, 71–94 (1974).

29. Kimura, K. & Kimura, A. Rab6 is required for the exocytosis of cortical granules and the recruitment of separase to the granules during the oocyte-to-embryo transition in *Caenorhabditis elegans*. J. Cell Sci. 125, 5897–5905 (2012).

30. Kondo, T. & Kimura, A. Choice between 1- and 2-furrow cytokinesis in *Caenorhabditis elegans* embryos with tripolar spindles. Mol. Biol. Cell 30, 2065–2075 (2019).

31. Johnston, W.L., Krizus, A. & Dennis, J.W. Eggshell chitin and chitin-interacting proteins prevent polyspermy in *C. elegans*. Curr. Biol. 20, 1932–1937 (2010).

32. Grant, B. & Hirsh, D. Receptor-mediated endocytosis in the *Caenorhabditis elegans* oocyte. Mol. Biol. Cell 10, 4311–4326 (1999).

