## Supplementary Information for "Cytoplasmic streaming drifts the polarity cue and specifies the cell polarity in *Caenorhabditis elegans* zygotes"

**Table S1.** ***C. elegans*** **strains used in this study.**

| Name | Genotype | Reference |
|---|---|---|
| CAL0182 | *fem-1(hc17) IV; unc-119(ed3) III, ruIs32 [pie-1p::GFP::H2B; unc-119(+)] III. ddIs6 [tbg-1::GFP; unc-119(+)] V.* | S1 |
| DE90 | *oxIs318 [spe-11p::mCherry::histone; unc-119(+)] II; unc-119(ed3 or e2498) ruIs32 III; ddIs6 V; dnIs17.* | S2 |
| DH1033 | *sqt-1(sc103) II; bIs1 X; bIs1[vit-2::GFP + rol-6(su1006)]* | S3 |
| CAL1041 | *unc-119(ed3) III; oxIs279 [pie-1p::GFP::H2B; unc-119(+)]., ltIs38[pAA1; pie-1::GFP::PH(PLC1delta1); unc-119(+)]* | This study |
| CAL0841 | *him-5(e1490) V; unc-119(ed3) III; oxIs318 II; unc-119(ed3 or e2498) ruIs32 III; ddIs6 V; dnIs17.* | This study |
| CAL1651 | *him-5(e1490) V; unc-119(ed3) III; oxIs318 II.* | This study |

References:


S1. Kondo T, Kimura A (2019) Choice between 1- and 2-furrow cytokinesis in *Caenorhabditis elegans* embryos with tripolar spindles. *Mol Biol Cell* 30:2065–2075.

S2. Johnston WL, Krizus A, Dennis JW (2010) Eggshell chitin and chitin-interacting proteins prevent polyspermy in *C. elegans*. *Curr Biol* 20:1932–1937.

S3. Grant B, Hirsh D (1999) Receptor-mediated endocytosis in the *Caenorhabditis elegans* oocyte. *Mol Biol Cell* 10:4311–4326.

### Supplementary Movie Legends

**Supplementary Movie 1**

Sperm pronucleus/centrosome complex (SPCC) dynamics of the PP-type zygote. White arrow indicates the position of SPCC. Time-lapse confocal fluorescence image of a zygote expressing GFP::histone, mCherry::histone, GFP::PH and GFP::TBG-1 was taken every 60 s. Eight different focal planes were stacked (Z-interval = 3 μm).

**Supplementary Movie 2**

Sperm pronucleus/centrosome complex (SPCC) dynamics of the DP-type zygote. White arrow indicates the position of SPCC. Time-lapse confocal fluorescence image of a zygote expressing GFP::histone, mCherry::histone, GFP::PH and GFP::TBG-1 was taken every 60 s. Eight different focal planes were stacked (Z-interval = 3 μm).

**Supplementary Movie 3**

Sperm pronucleus/centrosome complex (SPCC) movement and the flow of yolk granules during MeiCS. White arrow indicates the position of SPCC. Time-lapse confocal fluorescence image of a zygote expressing GFP::VIT-2 and mCherry::histone was taken every 5 s.
